# Decoding assembly of alpha-helical transmembrane pores through intermediate states

**DOI:** 10.1101/2021.09.08.459409

**Authors:** Neethu Puthumadathil, Greeshma S Nair, Smrithi R Krishnan, Kozhinjampara R Mahendran

**Affiliations:** Membrane Biology Laboratory, Transdisciplinary Research Program, Rajiv Gandhi Centre for Biotechnology, Thiruvananthapuram 695014, India; Manipal Academy of Higher Education, Manipal, Karnataka, India-576104

**Keywords:** Alpha-helical, pores, membrane, conductance, intermediate states, single-channel

## Abstract

Membrane-active pore-forming alpha-helical peptides and proteins are well known for their dynamic assembly mechanism and it has been critical to delineate the pore-forming structures in the membrane. Previously, attempts have been made to elucidate their assembly mechanism and there is a large gap due to complex pathways by which these membrane-active pores impart their effect. Here we demonstrate the multi-step structural assembly pathway of alpha-helical peptide pores formed by a 37 amino-acid synthetic peptide, pPorU based on the natural porin from *Corynebacterium urealyticum* using single-channel electrical recordings. More specifically, we report detectable intermediates states during membrane insertion and pore formation of pPorU. The fully assembled pore is functional and exhibited unusually large stable conductance and voltage-dependent gating, generally applicable to a range of pore-forming proteins. Furthermore, we used rationally designed mutants to understand the role of specific amino acids in the assembly of these peptide pores. Mutant peptides that differ from wild-type peptides produced noisy, unstable intermediate states and low conductance pores, demonstrating sequence specificity in the pore-formation process supported by molecular dynamics simulations. We suggest that our study contributes to understanding the mechanism of action of alpha-helical pores and antimicrobial peptides and should be of broad interest to bioengineers to build peptide-based nanopore sensors.

## Introduction

Membrane proteins carry out an extensive range of biological functions, including transport of ions, nutrients and metabolites, signal transduction, cell adhesion, anchoring other proteins, organizing organelles and cell shape.^1, 2^ There are two main structural classes of membrane proteins: alpha-helix and β barrels, where alpha-helices are more abundant than β-barrels.^1^ One such class of membrane proteins, i.e., pore-forming toxins (PFTs), are essential virulence factors secreted as soluble monomers that assemble into a structured oligomeric functional pore at the target membranes.^3-7^ Their mechanism of action involves the conversion of a prepore state to a pore via a complex assembly pathway (comprising/consisting of) transient intermediate steps.^3, 8-11^ Importantly, understanding their discrete dynamic assembly mechanism is vital in the development of therapeutics.^7^ Various techniques have been used to understand the assembly pathway of PFTs, including X-Ray crystallography, cryo-EM and fast-scan atomic force microscopy.^3, 7, 12, 13^ Interestingly, PFTs undergo large conformational changes depending on their native environment and hence, it is important to use techniques that provide information on the assembly dynamics in real-time in the membrane.^3, 9^

Notably, single-channel electrical recording is a method of choice for the characterization of pore-forming proteins as it provides insights into their structural and functional properties in the membrane environment.^9, 14^ Notably, the prepore state for pore-forming proteins is generally electrically silent with non-conducting states, and these states are most likely short-lived. ^8, 9, 15^ In addition to their potential in the development of therapeutics, certain PFTs such as α-hemolysin, ClyA, FraC, aerolysin have been used as nanopore sensors due to their unique architecture and pore geometry.^16-23^ Therefore, building sophisticated large-diameter pores are appropriate for large analytes such as nucleic acids, proteins and polysaccharides.^24^ Remarkably, there is great interest in developing pores formed from short synthetic peptides. Alpha-helical assemblies are particularly exciting targets due to their similarities to natural ion channels and pores based on alpha-helices are little-known.^25-29^ Notably, there are many obstacles in understanding alpha-helical assemblies due to their complex refolding mechanism and non-specific interactions with the membrane.^30-32^ Previously, we assembled a large synthetic transmembrane pore from 40-amino-acid alpha-helical peptides.^33^ The pore is charge selective, functional and capable of conducting ions and binding blockers such as cyclic oligosaccharides.^33^ We demonstrated charge-selective translocation of differently charged peptides through these synthetic alpha-helical pores.^34^

Here, we report a synthetic transmembrane pore, pPorU, built from 37-amino-acid peptides, corresponding to natural pore PorACur derived from *Corynebacterium urealyticum*.^35^ There were certain unique features to pPorU, including a tryptophan residue in the middle of the sequence compared to the terminal in other alpha-helical transmembrane motifs, proline and a cluster of phenylalanine residues resulting in further interest in the pore **(Fig 1a and Sfig 1)**. We probed the structure, assembly and conductance properties of these channels using a combination of peptide redesign and synthesis, solution-phase biophysics and single-channel recordings. We observed well-defined intermediate states during membrane insertion and pore formation of amphipathic alpha-helices. We suggest these synthetic peptides can act as a template to provide a model for understanding the assembly pathway of large pores, such as that found in bacterial toxins, pore-forming proteins of the immune system and antimicrobial peptides.^3, 36-38^ Our findings have important implications in biotechnology for designing pores for applications in nanopore technology, single-molecule nanopore chemistry and synthetic biology.

**Figure 1:**
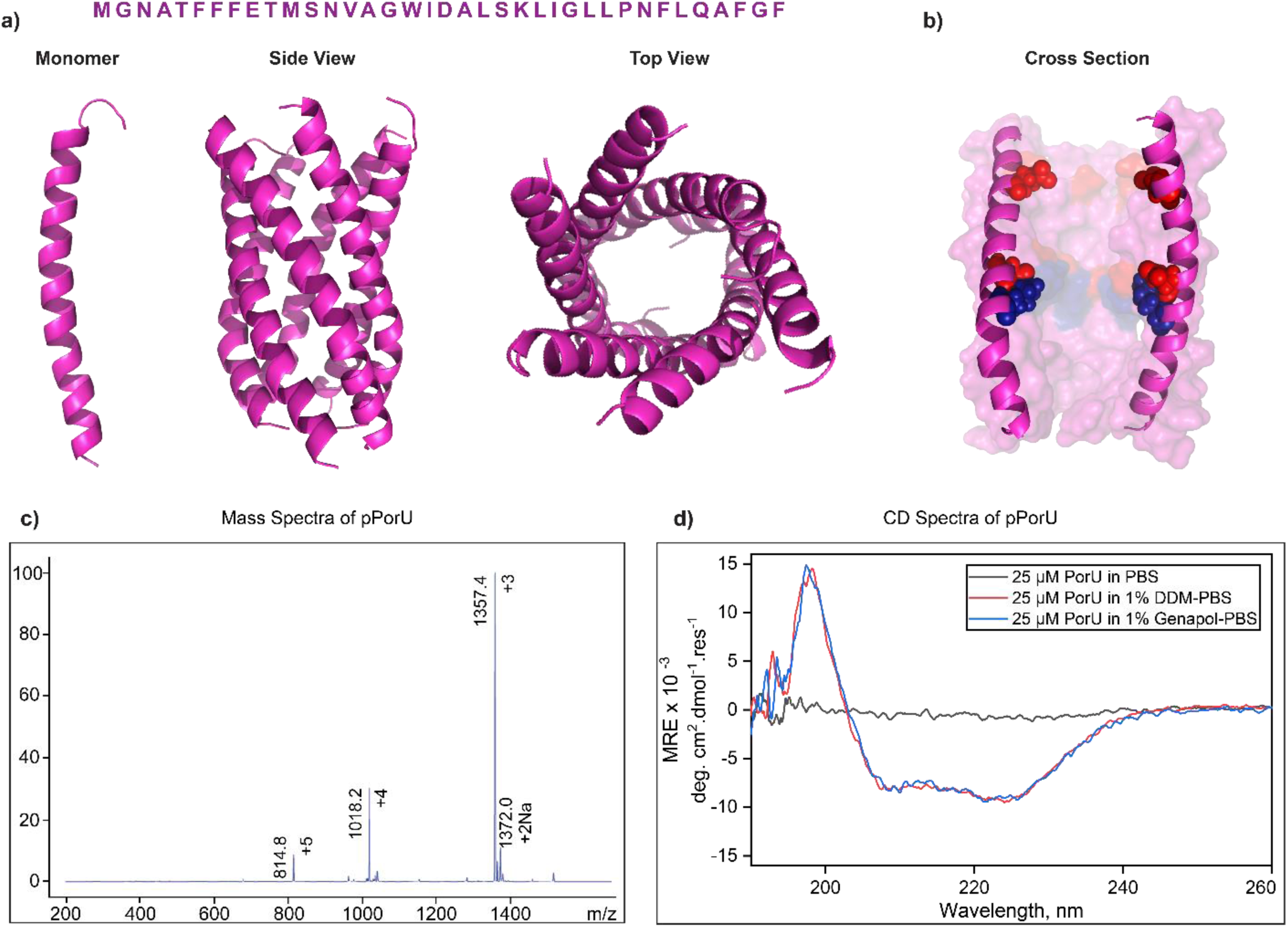
Biophysical properties of pPorU peptides. **a)** Modeled structure (Side view and Top view) and sequence of pPorU. **b)** Cross-sectional view of pPorU with positive residues in blue and negative residues in red. **c)** Mass spectra of pPorU peptide **d)** CD spectra of 25 µM pPorU in PBS (black), 1% DDM (red) and 1% Genapol (blue).

## Results and Discussion

### Biophysical properties of pPorU peptides

The 37 amino acid pPorU peptide was designed based on the sequence of natural porin PorACur derived from *Corynebacterium urealyticum* and synthesized by solid-phase synthesis **(Fig 1a)**.

The modeled structure (**Fig 1b and Sfig 1**) shows the cross-section of the pore with its electrostatic distribution (positively charged residue in blue and negatively charged residues in red). Both positively charged residues and denser negatively charged residues line the lumen of the pore, which might act as a binding site for charged analytes and contribute to the charge selectivity of the pore. The peptides were purified by reverse-phase high-performance liquid chromatography (HPLC) and the mass of the peptide was confirmed using mass spectroscopy (**Fig 1c**). Next, we investigated the secondary structure of the peptide using circular dichroism (CD) spectroscopy. A 25 µM peptide solution was prepared in different detergents such as 1% n-dodecyl-B-D-maltoside (DDM) and 1% Genapol solubilized in phosphate-buffered saline (PBS) (**Fig 1d**). The peptide exists in random coil conformation in PBS and folds itself into an alpha-helix conformation in the presence of detergent micelles **(Fig 1d)**.

### Electrical properties of pPorU peptides

The peptide (pPorU) was solubilized in 0.1% DDM and introduced into 1,2-diphytanoyl-sn-glycero-3-phosphocholine (DPhPC) planar lipid bilayer for characterization using single-channel electrical recordings in electrolyte buffer (1 M KCl, 10 mM HEPES, pH 7.4). The insertion of pPorU peptides into DPhPC bilayers was studied at different voltages ranging from +50 mV to +270 mV **(Fig 2a, Sfig 2 and Sfig 3)**. Remarkably, at a very high potential difference (≥ +200 mV), the pPorU peptide formed a large pore with a steady and stable conductance state in the lipid bilayers. Interestingly, we identified intermediates in pore assembly during peptide insertion: a small intermediate state (I1) and a large intermediate conductance state (I2). Finally, these two intermediate steps formed a large stable pore (L) (**Fig 2, Fig 3 and Sfig 2**). The timeline for forming large stable pores from intermediates ranges from milliseconds to several seconds, precisely resolved by single-channel recordings **(Sfig 3)**.

**Figure 2:**
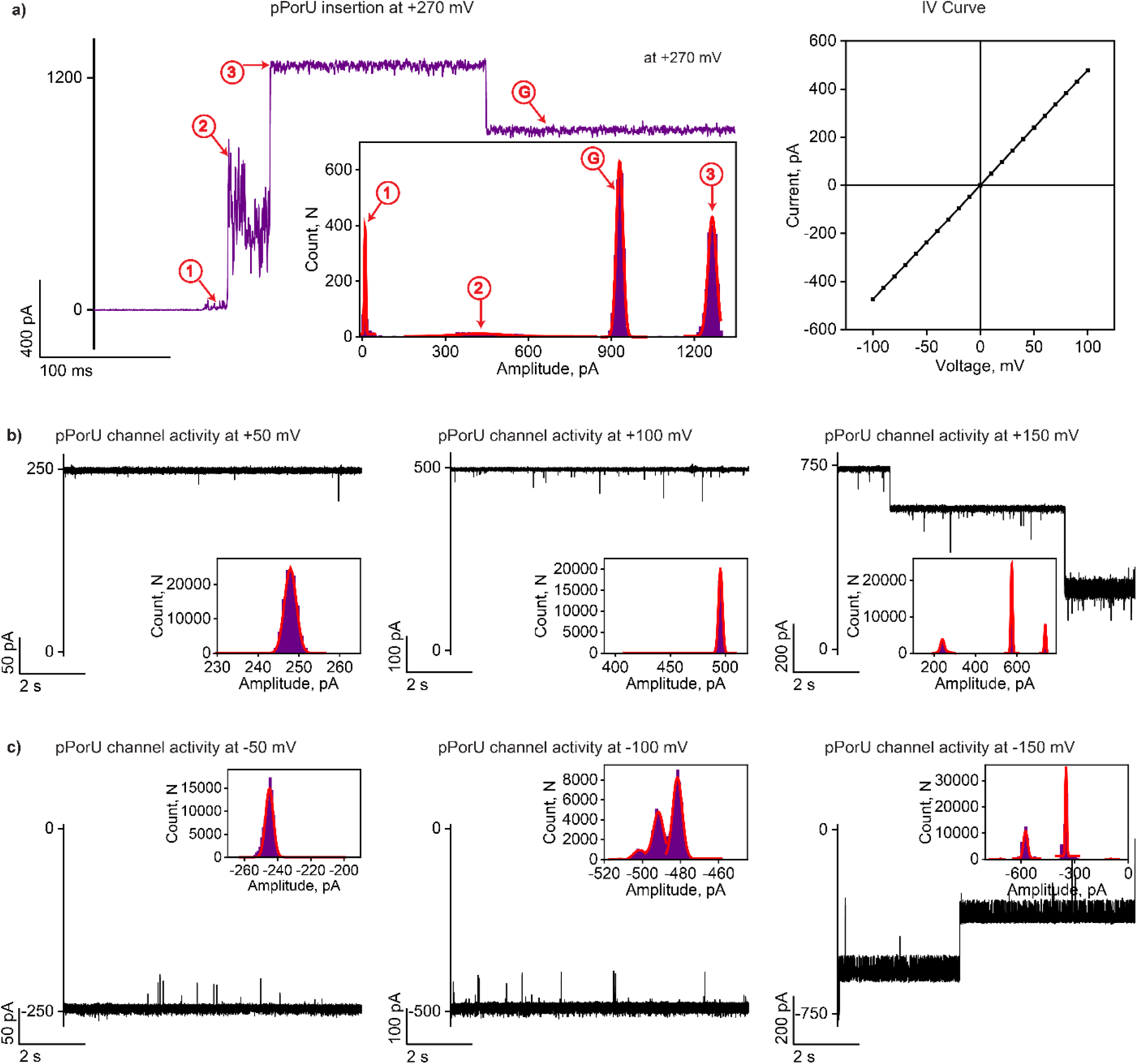
Electrical properties of pPorU pore. **a)** Electrical recording of single pPorU channel insertion into DPhPC lipid bilayer at +270 mV and the corresponding current amplitude histogram. Current-voltage curve is shown. **b)** Electrical recordings of single pPorU pore at +50 mV, **+**100 mV and +150 mV. **c)** Electrical recordings of single pPorU pore at -50 mV, **-**100 mV and -150 mV. The current signals were filtered at 2 kHz and sampled at 10 kHz. Electrolyte: 1 M KCl, 10 mM HEPES, pH 7.4.

**Figure 3:**
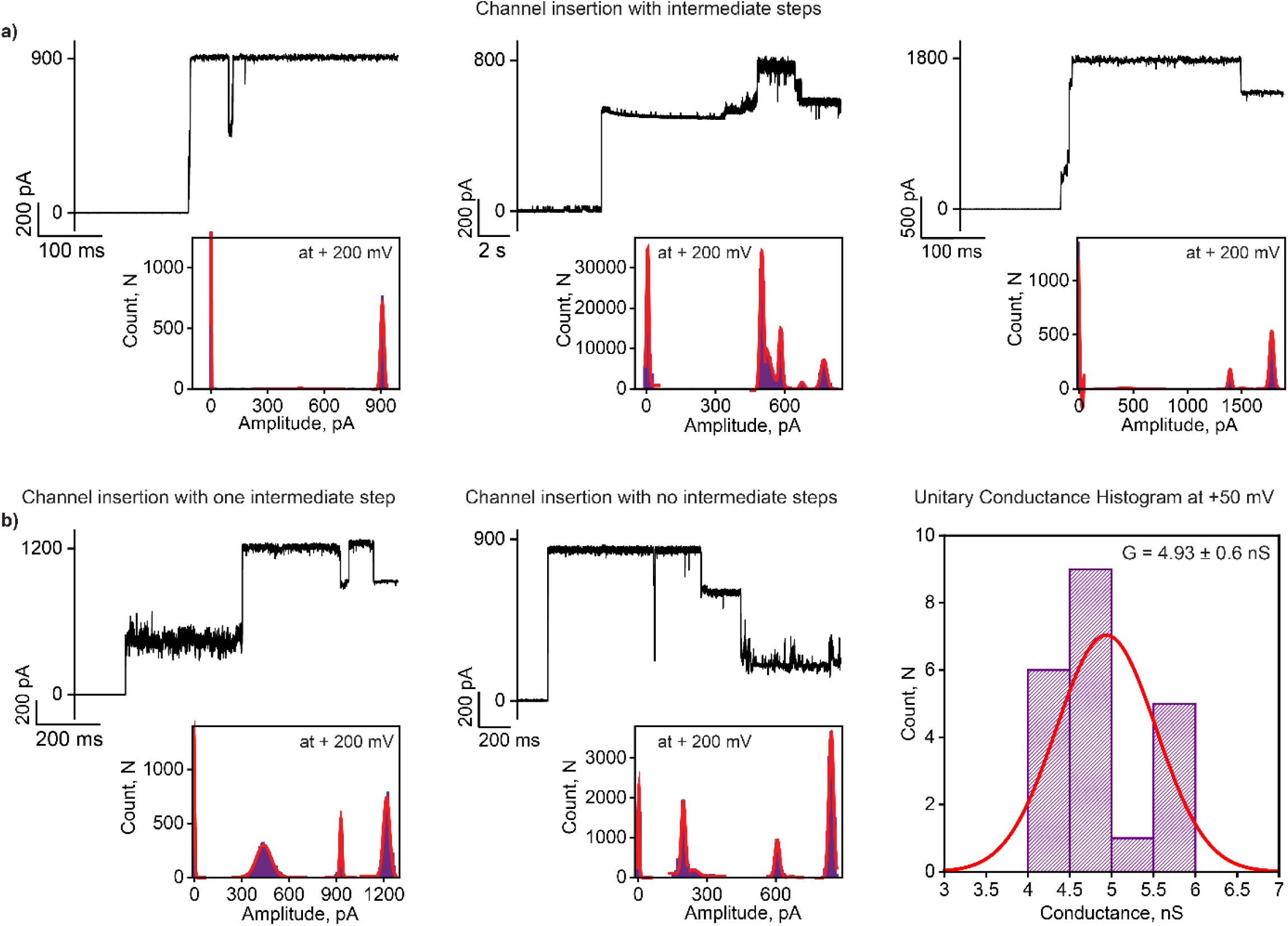
Electrical properties of pPorU pore. **a)** Electrical recording of pPorU channel insertions with intermediate steps into DPhPC lipid bilayer at +200 mV and the corresponding current-amplitude histogram. **b)** Electrical recording of pPorU channel insertions with and without intermediate steps into DPhPC lipid bilayer at +200 mV and the corresponding current-amplitude histogram. The unitary conductance histogram at +50 mV was obtained by fitting the distribution to a single Gaussian (n = 16). The current signals were filtered at 2 kHz and sampled at 10 kHz. Electrolyte: 1 M KCl, 10 mM HEPES, pH 7.4.

Following this, several insertions of pPorU peptides at very high voltages were recorded. The transition into the large pore via the two states was observed multiple times (**Fig. 2a** and **Sfig 2**). Notably, the final conductance state remains in an open state at voltages up to ± 100 mV (**Fig. 2b and Sfig 3)**. At higher voltages (≥ ±150 mV), gating was observed and the pore fluctuated between open and closed conductance states **(Fig 2b, Fig 2c and Sfig 4)**. Voltage gating is an intrinsic property observed in most pores, with the threshold potential of gating varies from pore to pore; however, the mechanism of gating is unknown. The first intermediate state (I1) showed transient ion current bursts and fluctuating current in the range of 2-5 pA, which varies from pore to pore (**Fig 2a, Sfig 2 and Sfig 3**). This state most likely represents the initial interaction of peptide monomers with the membrane, which consecutively drives the oligomerization of peptides. This state has a time duration in milliseconds and could represent a state before transmembrane pore formation, characteristic of a prepore state **(Fig 3a and 3b)**. Notably, the I1 state opened into an intermediate noisy conductance state (I2), which exhibited slight current fluctuations, and this distinct conductance state was calculated to be ∼2 nS derived from the all points amplitude histogram **(Fig 2a, Fig 3, 3b, Sfig 2 and Sfig 3)**. This state mostly lasts for milliseconds which could represent the pore continuing oligomerization in the membrane environment. Occasionally, we observed large pore formation devoid of one or both the intermediate steps **(Fig 3b)**. Most of the time, the assembly of the large stable pore is driven by unsteady intermediate conductance states, specifically at high voltages (± 200 mV) **(Sfig 3)**. The conductance histogram was obtained based on the channel insertions and a unitary conductance was calculated to be 4.93 ± 0.6 nS (n=16), indicating the formation of uniform pores of a single oligomerization state **(Fig 3b and Sfig 3)**. Based on this data, we propose that the larger conductance pore formed by pPorU peptides can be used as a nanopore sensor for single-molecule sensing of analytes **(Fig 4 and Sfig 5)**.

**Figure 4:**
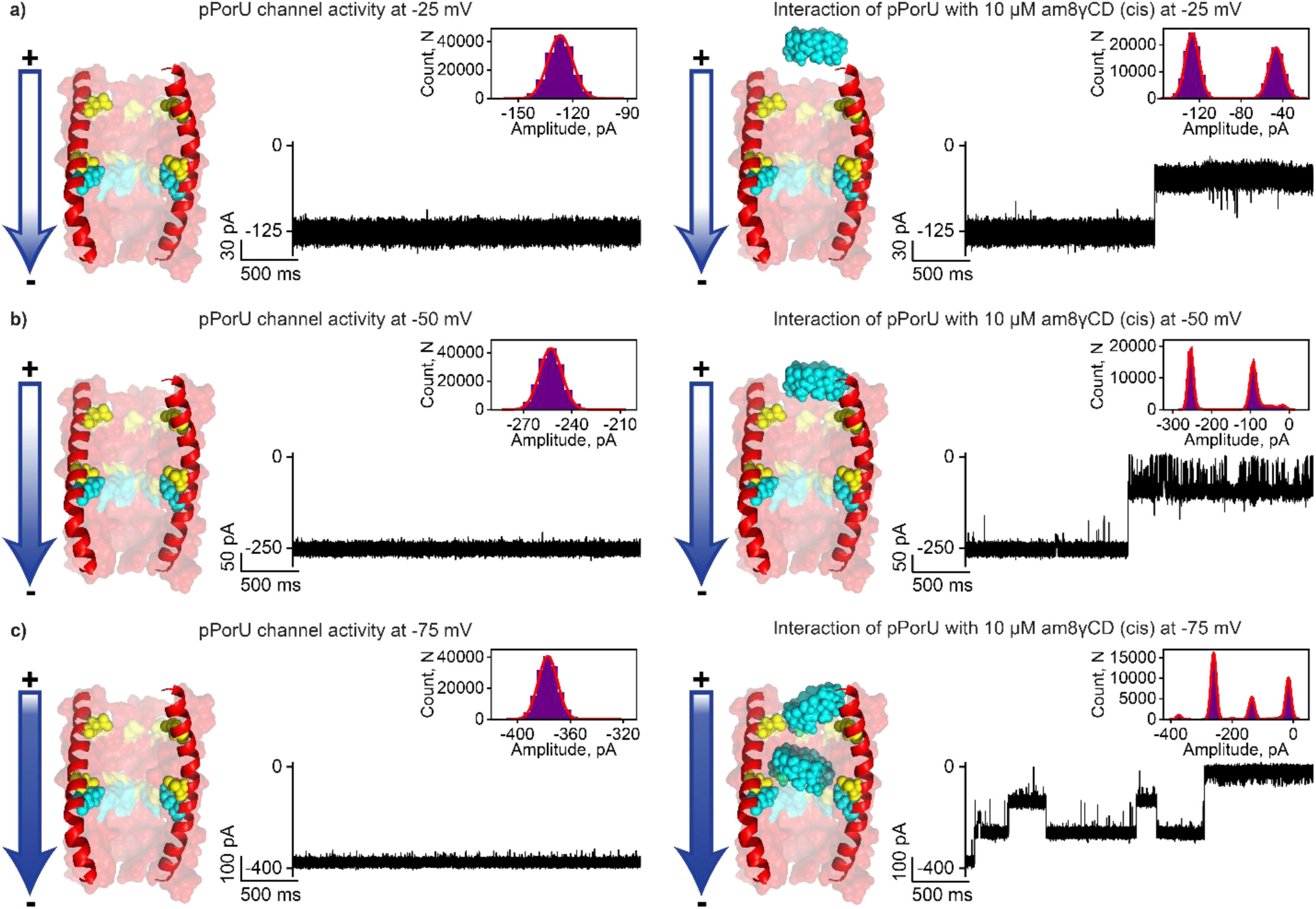
Interaction of pPorU pore with cationic cyclodextrin. **a)** Electrical recordings showing single pPorU pore at -25 mV and the interaction of cationic gamma-cyclodextrin (am8γCD) with single pPorU pore (10 μM, cis) at -25mV. **b)** Electrical recordings showing single pPorU pore at -50 mV and the interaction of cationic gamma-cyclodextrin (am8γCD) with single pPorU pore (10 μM, cis) at -50 mV. **c)** Electrical recordings showing single pPorU pore at -75 mV and the interaction of cationic gamma-cyclodextrin (am8γCD) with single pPorU pore (10 μM, cis) at -75 mV (n=2). The corresponding amplitude histogram is shown in the inset. Current signals were filtered at 10 kHz and sampled at 50 kHz. Electrolyte: 1 M KCl, 10 mM HEPES, pH 7.4.

The modeled structure of pPorU revealed that the pore lumen is lined by negatively charged residues which can serve as a binding site for large cationic molecules. Accordingly, we examined the interaction of large cationic octasaccharide am8γCD with pPorU (L state) by measuring the ion current blocking at different voltages to elucidate the functionality of the pore. (**Fig 4 and Sfig 5)**. The interaction of am8γCD with the pPorU L state was studied at the voltages where the pore remained fully open. This allowed us to distinguish between CD-induced blocking and voltage gating **(Fig 4 and Sfig 5)**. The addition of am8γCD (10 µM) to the cis side resulted in ionic current blockages only at negative voltages as CDs are driven through the pore by the applied electric field **(Fig 4a-4c and Sfig 5)**. At -25 mV, the blockage events are infrequent, and the number of ion current blockage events increased with an increase in the voltage at -50 mV **(Fig 4a-4c and Sfig 5)**.

At -75 mV, am8γCD blocked the pore in several steps of decreasing and increasing conductance, indicating multiple CD binding sites in the pore **(Fig 4c and Sfig 5)**. We propose that the negative voltage electrophoretically pulls cationic CDs into the pore and binds to the negatively charged residues. As expected at positive voltages, we observed no blockage events confirming voltage-dependent electrostatic charge-based interaction of CDs with the pore (**Sfig 5)**. The insertion of pPorU peptides into planar lipid membranes was studied at lower voltages (±50 mV and ±100 mV) and surprisingly, large-conductance pore formation was not detected irrespective of different concentrations of the pPorU peptides added into the bilayer chamber **(Fig 5a-5c and Sfig 6)**.

**Figure 5:**
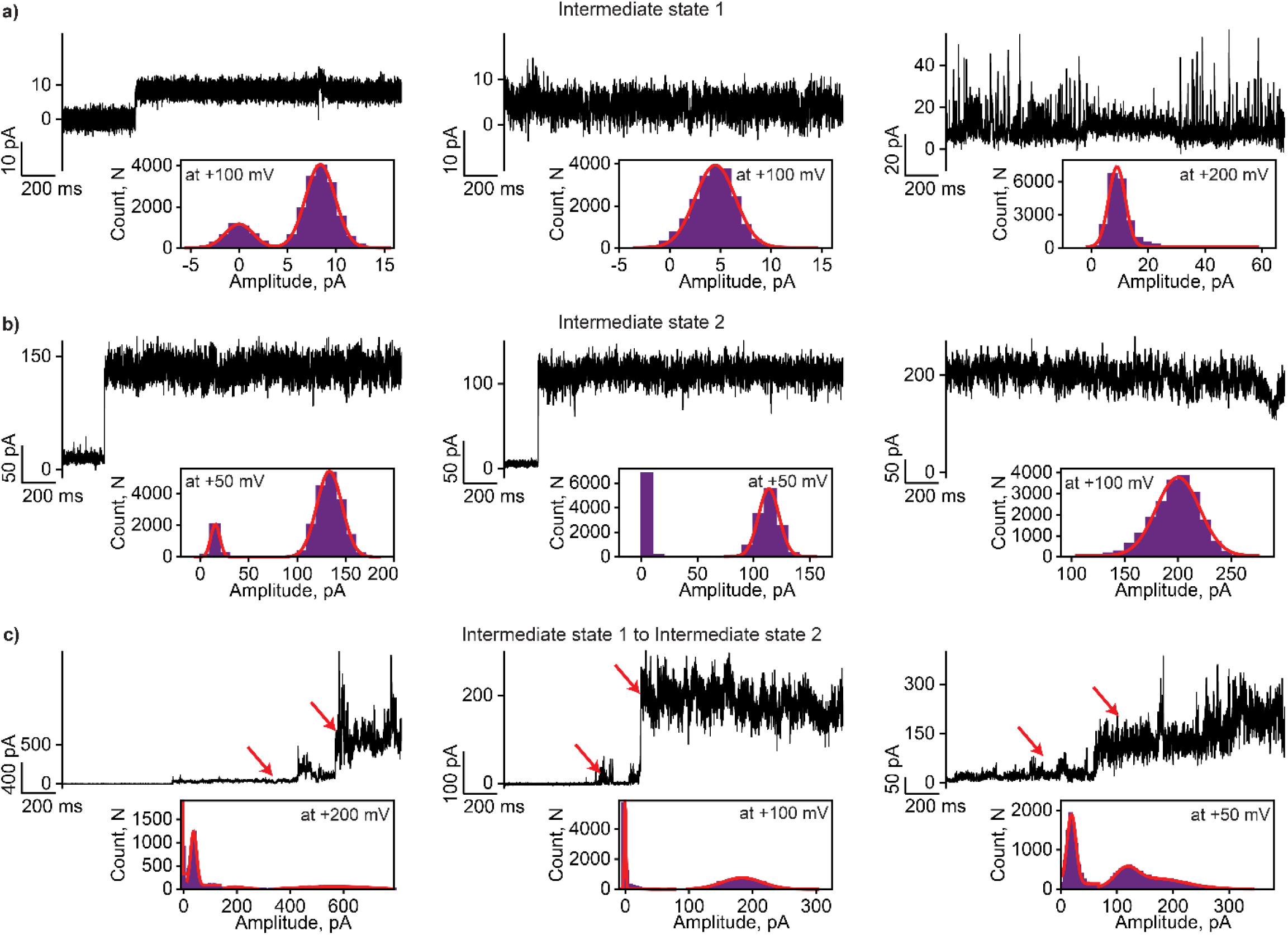
Electrical properties of pPorU pore. **a)** Electrical recording of intermediate conductance states at lower voltages showing low conductance states. **b)** Electrical recording of different intermediate states at lower voltages showing higher conductance states. **c)** Electrical recording of intermediate states at lower voltages showing the transition of different conductance states. The corresponding amplitude histogram is shown in the inset. Current signals were filtered at 2 kHz and sampled at 10 kHz. Electrolyte: 1 M KCl, 10 mM HEPES, pH 7.4.

At lower voltages, pPorU peptides formed pores of varying conductance states, indicating heterogeneous forms in the membrane **(Fig 5a-5c and Sfig 6)**. We observed that peptides formed transient spikes and fluctuating low conductance pores (2-10 pA). In some instances, the peptides formed noisy high conductance pores (2 nS), which fluctuated into lower conductance states **(Fig 5b and Sfig 6)**. We show that the pPorU peptide interaction with the membrane could be trapped in the intermediates steps by applying lower potential during pore insertion, allowing characterization of the assembly pathway. This indicates that the pPorU peptides are not assembled into stable oligomers in the membrane at lower voltages and pores are trapped in the intermediate state conformation **(Fig 5 and Sfig 6)**. In comparison, higher voltages induce the formation of a large stable pore in the membrane via intermediates.

### The specificity of pore assembly: pPorU mutants

To elucidate the pore assembly pathway based on the time-resolved intermediate states detected during pPorU peptide insertions, we rationally designed mutant peptides based on the modeled pPorU. We specifically designed two mutants, pPorU-W17C and pPorU-P29C, to understand the role of these amino acids in the assembly of alpha-helical peptide pores (**Fig 6, Sfig 7, and Sfig 8**). Tryptophan residues, present in a higher concentration in membrane proteins than soluble proteins, have been predicted to play an important role in anchoring pores in the membrane.^39, 40^ Further, the presence of tryptophan in the middle of the pPorU sequence in contrast to the terminal position as located in other membrane proteins, including antimicrobial peptides, stimulated us to focus on this particular amino acid.

**Figure 6:**
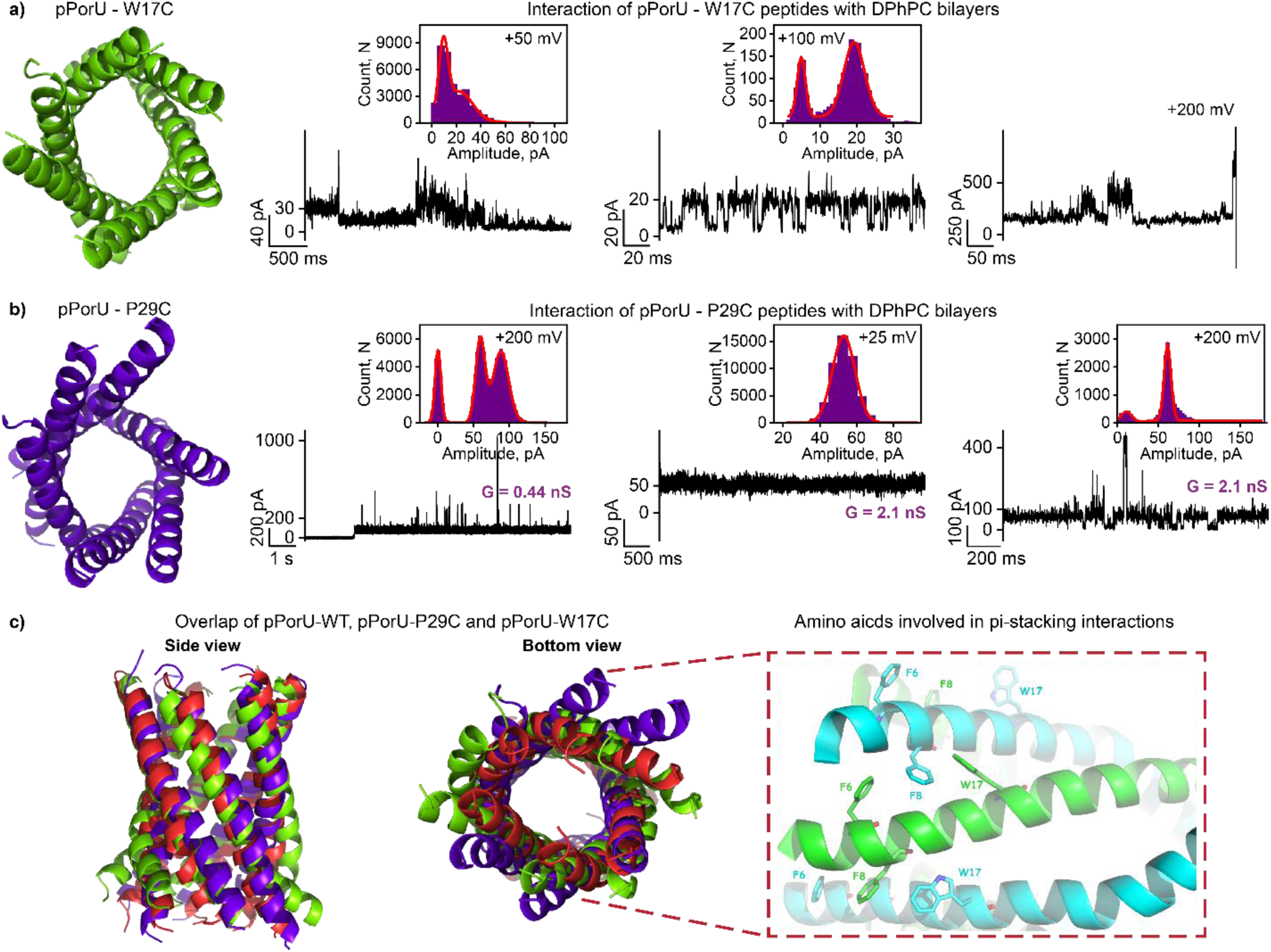
pPorU mutants and molecular dynamics simulations of pPorU pores. **a)** Modeled structure of pPorU-W17C. Electrical recording of pPorU-W17C peptides showing fluctuations at +50 mV. Electrical recording of the pPorU-W17C peptides showing two states at +100 mV and bilayer disruption at +200 mV. **b)** Modeled structure of pPorU-P29C. Electrical recordings of pPorU-P29C showing low conductance channel activity at +200 mV and +25 mV. The corresponding amplitude histogram is shown in the inset. Current signals were filtered at 2 kHz and sampled at 10 kHz. Electrolyte: 1 M KCl, 10 mM HEPES, pH 7.4. **c)** Modelled hexameric pPorU-WT overlapped with pPorU-W17C (green) and pPorU-P29C (blue). Side view of the WT pPorU (red) structure, showing critical residues involved in a regular series of pi-stacking interactions that stabilize the N-terminal portion of the pore (Right).

The second residue of interest, proline, has been known to form kinks in the secondary structure of peptides and proteins.^41-43^ Previous studies have shown that proline residues in membrane proteins (transmembrane alpha-helices) induce distortions and play a key role in helix packing, thus contributing to dynamic flexibility.^44-47^ The peptides pPorU-W17C and pPorU-P29C were synthesized by solid-phase synthesis and purified similar to WT pPorU by HPLC and the mass of the peptides was confirmed using mass spectroscopy **(Sfig7 and Sfig 8)**. Accordingly, we examined the pore-forming properties of the mutant peptides using single-channel electrical recordings. The insertion of pPorU-W17C peptides was studied at lower and higher voltages (+200 mV) **(Fig 6a and Sfig 7)**. The mutant pPorU-W17C (n=27) formed fluctuating channels of varying conductance in the membrane, indicating non-uniform unstable assemblies **(Fig 6a and Sfig 7)**. Furthermore, the fluctuation increased with increasing voltages, with the ion conductance ranging from 0.05 nS to 2 nS **(Fig. 6a and Sfig 7)**. In some instances, the peptide formed transient pores with ion current spikes of short duration and low amplitude (<+50 pA). Notably, at higher voltages (+200 mV), we observed a drop in the ion current to almost zero (n=9), most likely indicative of the unstable peptide oligomers leaving the bilayer due to dissociation **(Fig. 6a and Sfig 7**). Based on these results, we hypothesize that due to the lack of anchorage provided by the tryptophan, the peptide could not insert into the bilayer in a stable conformation to form well-defined pores. We suggest that tryptophan mutant non-specifically ruptures the membrane-like antimicrobial peptides.

Unlike the tryptophan mutant, the proline mutant pPorU-P29C (**Fig. 6b and Sfig 8**) inserted into the membrane rapidly, emphasizing the importance of tryptophan in membrane insertion and anchorage. At +200 mV, pPorU-P29C peptides formed pores in the membrane (n=20) with two distinct channel properties. The first pore type produced a very low current state (less than 10 pA) with intermittent transient noisy spikes, most likely indicating the initial binding of peptides with the membrane. This particular state is similar to the prepore state or first intermediate state exhibited by the WT pPorU (**Fig. 6b and Sfig 8**). The second type pore produced a slightly fluctuating state with an increased conductance of ∼2 nS, similar to the second intermediate state formed by WT pPorU. The pore produced noisy conductance states in some instances, although a stable conductance state is generally observed. In contrast to the tryptophan mutant, the proline mutant remained in the membrane for longer durations, demonstrating considerably stable oligomer formation in the membrane environment (**Fig. 6b and Sfig 8**). Interestingly, the proline mutant did not form the large stable conductance channels (5 nS in 1M KCl) as seen for WT pPorU. We suggest that this particular mutant is trapped in the intermediate steps indicating partial oligomer conformation formation. Thus, proline could be playing a role in the transition of the peptide monomers and oligomers to the final stable conformation through the different intermediate states. Importantly previous studies support the role of proline in controlling the flexibility of alpha-helical peptides in the membrane environment.^45-47^ Remarkably, the drastic difference in the single-channel electrical and biophysical properties of WT and pPorU mutants demonstrates the sequence specificity of alpha-helical peptide pores. The major change in alpha-helical pore’s structural and functional properties with a single amino-acid substitution affirms the exciting features of alpha-helical pore assemblies. Accordingly, the flexibility and tunability of alpha-helices associated with balancing their self-assembly and membrane solubility are essential for assembling defined structures.

### Molecular Dynamics Simulations

So far, we have elucidated the assembly pathway of the peptide pore pPorU using single-channel electrical recordings supported by the rationally designed mutants. We used molecular dynamics simulations to address why small sequence changes make a difference in single-channel properties and subsequently investigate the specificity in forming the large pores. Through the extent of the equilibrations and simulations, the WT pPorU and two mutational variant pores sustained semi-regular hexagonal arrangements, with the most stable feature in all three structures being a well-conserved network of inter-peptide salt bridges between Asp 19 and Lys 23 of neighboring peptides. Beyond this, however, MD simulations indicate significant differences among the propensity of the different variants to sustain coherent pore-like structures in the vicinity of the pore mouths **(Fig 6c and Sfig 9)**. Trp 17 proves to be a crucial determinant of pore viability and pPorU W17C pore structure tends to degenerate more irregularly. This destabilization illustrates the stable role of Trp 17 in anchoring a series of pi-stacking arrangements that are crucial to preserving a regular series of inter-peptide interactions within the N-terminal region of the pore **(Fig 6c and Sfig 9)**. Specifically, Trp 17 on one peptide generally has a favorable interaction with a Phe 8 of a neighboring peptide, which, in turn, pi-stacks with Phe 6 on the first peptide. Substituting Cysteine in place of Trp 17 negates the inter-peptide coupling with Phe 8, consequently, ceases to stabilize Phe 6. The loss of these contacts produces a general lack of pore integrity for the pPorU W17C peptide.

Nevertheless, the significant lipophilicity inherent in the pPorU W17C peptide likely ensures that it will interact with cell membranes, but this is unlikely to translate into the formation of stable membrane-spanning pores (**Fig. 6a, 6c and Sfig 9)**. Notably, pPorU P29C pore exhibits substantial loosening of inter-peptide contacts in the C-terminal region of the pore. Notably, the native form sustains a modest number of inter-peptide H-bonds within the C-terminal region via associations between Asn 30 and Gln 33. Therefore, the mutation of Pro 29 to Cysteine disrupts nearly all of these C-terminal contacts, replacing the inter-peptide H-bonds with intra-peptide coupling **(Fig. 6b, 6c and Sfig 9)**. Without these pore-stabilizing features, the C-terminal segments display amphiphilicity, with appreciable lipophilic surfaces that tend to maximize contacts with lipids. This leads to a pPorU P29C pore that fans out into a broader structure lacking the barrel shape of a stable membrane pore. The importance of Pro 29 in sustaining inter-peptide H-bonds is interesting since proline does not, in itself, offer opportunities for side-chain H-bonding coupling **(Fig. 6b, 6c and Sfig 9)**. Proline is valuable for pore stability because it kinks the peptide helix in a manner that induces the C-terminus of one peptide to more closely approach one of its immediately adjacent peptide neighbors, thus favoring inter-peptide N30 and Q33 H-bonds over intra-peptide associations.

## Discussion

The natural porin PorACur derived from *Corynebacterium urealyticum* is a membrane-spanning alpha-helical pore, and our goal was to see if a synthetic peptide based on this natural porin can spontaneously self-assemble into a pore in the membrane environment.^35^ The 37-amino-acid peptide, pPorU, was found to form a pore and details of its structure and the assembly pathway were obtained. Our model for the pore assembly based on electrical recordings is that individual pPorU peptide monomers insert into the bilayer to form a small intermediate (I1) state, revealing membrane association of the peptides. Further, the small intermediate I1 then assemble spontaneously in the membrane to form the large intermediate state (I2), which showed a noisy well-defined conductance state indicating the formation of the incomplete pores. Finally, (I2) state transforms into a fully opened stable conductance state, revealing the assembly of complete stable pores (L), which remained in the lipid membrane in the same conformation for longer durations **(Sfig 10)**. Remarkably, we show that this particular model of stable pore assembly is limited to high voltages, whereas unstable noisy intermediate state pores are formed at lower voltages. We also comment on the potential role of specific amino acids in the pore’s transition via different intermediates to the large conductance pore. The C-terminal D4 domain of the 340-kDa Wza polysaccharide transporters is an octameric alpha-helix barrel that spans the outer membrane of various Gram-negative bacteria.^28^ Previous studies have shown that 35-amino acid, synthetic D4 peptides corresponding to the components of this domain insert into lipid bilayers to form pores via precursor state, low conductance and high conductance state. Interestingly, in contrast to pPorU pores, the cWza pore transitions were reversible.^28^ pPorU also exhibited unusually high conductance of ∼5.0 nS, whereas fully open cWza exhibited ∼0.9 nS conductance and did not remain in the bilayer as oligomers disassemble to monomeric peptide conformations. Based on this, we suggest that the intermediate steps observed during pPorU pores could represent membrane binding, different oligomer conformations and pore states proposed in the assembly of PFTs.^7^ This understanding could provide a better understanding of their assembly pathway and reveal amino-acid targets that could specifically hamper, binding or transition of the PFTs, trapping them in their non-toxic states. This could also lead to the development of more effective therapeutics against bacteria that produce membrane attacking PFTs.^7^

Previously, we have shown that pPorA self-assembled as an octamer in the SDS PAGE and gel extracted octamer readily inserted into planar lipid bilayers and formed stable cation-selective pores.^33, 34, 48^ In contrast to pPorA, the pPorU peptides run on an SDS-PAGE gel formed only monomers, indicating the absence of self-assembled stable oligomers.^33^ This drastic difference in the assembly mechanism of pPorU and pPorA demonstrates the sequence specificity of alpha-helical peptide pore formation in planar lipid bilayers. Similar to pPorA, the pPorU peptides formed cation-selective pores that were blocked by cationic cyclic oligosaccharides indicating the presence of predominant negatively charged residues similar to pPorA. Additionally, we suggest that the presence of a tryptophan residue in the middle of the pPorU sequence along with proline and a cluster of phenylalanine residues gives specificity in the formation of large-conductance pores. We also demonstrate their potential as a nanopore sensor for sensing complex macromolecules as the demand for developing superior pores continues. For instance, these large-diameter pores can be applied for single-molecule sensing of large analytes, including antibodies, complex nucleic acids and polysaccharides.^25, 49^

## Conclusion

Understanding the molecular assembly mechanism of membrane-active peptides and proteins is very challenging. We characterized the assembly of short alpha-helical peptide pores based on naturally occurring alpha-helical porins using single-channel electrical recordings. We elucidated the intermediate steps involved in the pore assembly from peptide monomers to a large pore oligomer in the lipid membrane. Furthermore, we characterized various mutants to decipher the critical amino acids’ role in the structural assembly pathway. We suggest that the understanding assembly mechanism of simple alpha-helical peptide pores sheds light on the mechanism of clinically relevant pore-forming proteins, including membrane-active toxins and antimicrobial peptides. This new class of alpha-helical pores showing very large-conductance can be used for nanopore technology and would allow the design of pores for biotechnology applications.

### Experimental section

#### Single-Channel Electrical Recordings

Planar lipid bilayer recordings were carried out by using bilayers of 1,2-diphytanoyl-snglycero-3-phosphocholine (DPhPC, Avanti Polar Lipids) formed across an aperture (∼100 μm in diameter) in a 25-μm thick polytetrafluoroethylene (Teflon) film (Goodfellow, Cambridge), which separated the apparatus into cis and trans compartments (600 μL each).^50^ Bilayers were formed by first pretreating the aperture with hexadecane in n-pentane (2 μL, 5%) on each side. Both compartments were then filled with the electrolyte solution (1 M KCl, 10 mM HEPES, pH 7.4), and DPhPC in n-pentane (2 μL, 5 mg mL^−1^) was added to both sides; after that, the solvent evaporated. A bilayer was formed when the electrolyte was raised, bringing the two lipid surface monolayers together at the aperture. The pPorA pores were formed by adding a peptide solution (cis side) in 0.1% DDM (0.1 μL, 25 μM) under an applied potential of +200 mV. DTT (1 μM) was added to the cis side to facilitate the insertion of cysteine mutants of pPorU peptides into the lipid bilayer. The cis compartment was connected to the grounded electrode, and the trans compartment was attached to the working electrode. A potential difference was applied through a pair of Ag/AgCl electrodes, set in 3% agarose containing 3.0 M KCl. The current was amplified using an Axopatch 200B amplifier, digitized with a Digidata 1550B, and recorded with pClamp 10.6 acquisition software (Molecular Devices, CA) with a low-pass filter frequency of 2 kHz and a sampling frequency of 10 kHz. pPorU blocking by cationic cyclodextrins was quantified using a statistical analysis of the pore in its unblocked and blocked states. These data were recorded with a low-pass filter frequency of 10 kHz and a sampling frequency of 50 kHz to resolve blocking events up to 100 μS. The data were analyzed and prepared for presentation with pClamp (version 10.6, Molecular Devices, CA) and Origin.

#### Circular Dichroism Spectroscopy

Circular dichroism spectra were obtained using Jasco J-810 spectropolarimeter. Peptide samples were prepared as 25 μM solutions in phosphate-buffered saline (PBS, 8.2 mM sodium phosphate, 1.8 mM potassium phosphate, 137 mM sodium chloride, and 2.7 mM potassium chloride at pH 7.4) with 1% DDM, 1% Genapol or 1% SDS. Spectra were collected using a 1 mm path length quartz cuvette at 20 °C. For each data set (in deg), baselines from the same buffer and cuvette were subtracted, and then data points were normalized for amide bond concentration and path length to give mean residue ellipticity (MRE; deg cm^2^ dmol^−1^ res^−1^).

#### Molecular Dynamics Simulations

Molecular dynamic simulations were performed to assess the conformational stability of two specific point mutations (W17C and P29C) of membrane-bound hexameric pPorU compared with the native pPorU pore. The pore structure’s initial approximation entailed a bundle of six regularly helical pPorU monomers assembled as an unstrained barrel of membrane-spanning length. From this initial structure, prospective structures for the W17C and P29C mutants were constructed by editing the native pPorU structure in PyMol using the mutagenesis wizard. Using the on-line Charmm GUI^51^, each resulting pore was embedded into a bilayer composed of POPC, POPE and POPG lipids in a 3:1:1 ratio, with cytoplasmic and extracellular buffers (extending 15.0 Å above and below the bilayer) composed of water, 67 K+ ions and 21 Cl-ions, thus comprising material and charge densities that reasonably approximate the dielectric within proximity to a POPC+POPE+POPG membrane. To derive a physically realistic starting point from which to assess the relative stability of the three pore models, extensive restrained constant pressure structural refinements were carried out in NAMD ^52^, representing the structures via the Charmm 3.6 force field.^53^ Given a thermostatic temperature of 303.15K, an initial 10,000 step relaxation was performed using quadratic positional restraints on all protein backbone atoms via a restraint scaling factor of 10.0. A second 12,500 step restraint relaxation was then conducted on the resulting system, using a scaling factor of 5.0. Four further relaxations were then conducted (of respective lengths 12,500, 25,000, 25,000, and 25,000 steps) with restraint scaling factors diminishing according to the progression 2.5, 1.0, 0.5 and 0.1). A final production simulation was then run for 4.0 nS, at constant pressure, with no positional restraints.

## Acknowledgments

This work was supported by the Ramalingaswami Re-entry fellowship of the Department of Biotechnology, Government of India (BT/RLF/Re-entry/49/2014). KRM acknowledges ‘Early Career Research Award’ of Science & Engineering Research Board (SERB), Department of Science and Technology, Government of India (ECR/2017/000533) and the research grant awarded by the Department of Biotechnology, Government of India (BT/PR34466/BRB/10/1830/2019) for supporting this work. NP is supported by the University Grants Commission, Government of India. SKR is supported by Senior Research Fellowship from the Indian Council of Medical Research (5/3/8/9/ITR-F/2020).

## Author contributions

NP, SKR and GN performed single-channel current recordings. NP analyzed the single-channel data and performed biophysical characterization of peptides. NP and KRM conceived the study, designed the experiments and wrote the paper.

## Supporting Information Available

Materials, text, and figures are available in the supplementary Information.

