## Supplementary file for "Decoding assembly of alpha-helical transmembrane pores through intermediate states"

**Supplementary Materials**

The following materials were used for the study: 1,2-diphytanoyl-*sn*-glycero-3-phosphocholine (DPhPC, Avanti Polar Lipids), pentane (Sigma-Aldrich Merck), hexadecane (Sigma-Aldrich Merck), n-dodecyl β-D-maltoside (DDM, Sigma-Aldrich Merck), Sodium dodecyl sulfate (SDS, Sigma-Aldrich), Genapol (Sigma-Aldrich), octakis-(6-amino-6-deoxy)-γ-cyclodextrin octahydrochloride (am$_8$γCD, AraChem Cyclodextrin-Shop), potassium chloride (Sigma-Aldrich Merck), 4-(2-hydroxyethyl)-1-piperazineethanesulfonic acid (HEPES, Sigma-Aldrich Merck), ethylenediaminetetraacetic acid disodium salt (EDTA, Sigma-Aldrich Merck), dithiothreitol (DTT, Sigma-Aldrich Merck), sodium chloride (Sigma-Aldrich Merck), 2-Propanol (Sigma-Aldrich Merck), Acetone(Sigma-Aldrich Merck), all other reagents (Sigma-Aldrich Merck). All pPorU peptides were purchased from Peptide Protein Research Ltd at >95% purity (HPLC) as lyophilized powders.

### Supplementary Figures

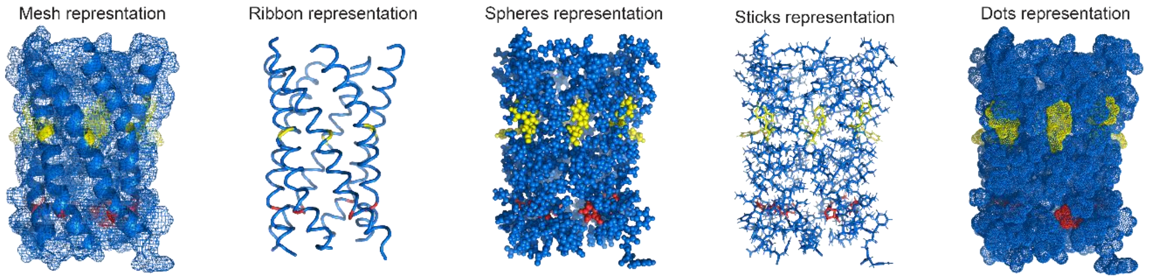


**Figure S1: Modeled Structure of pPorU pores**

Modeled structure of pPorU pores in different representations: Mesh, Ribbon, Spheres, Sticks and Dots. The tryptophan residues are highlighted in yellow and proline residues are highlighted in red.

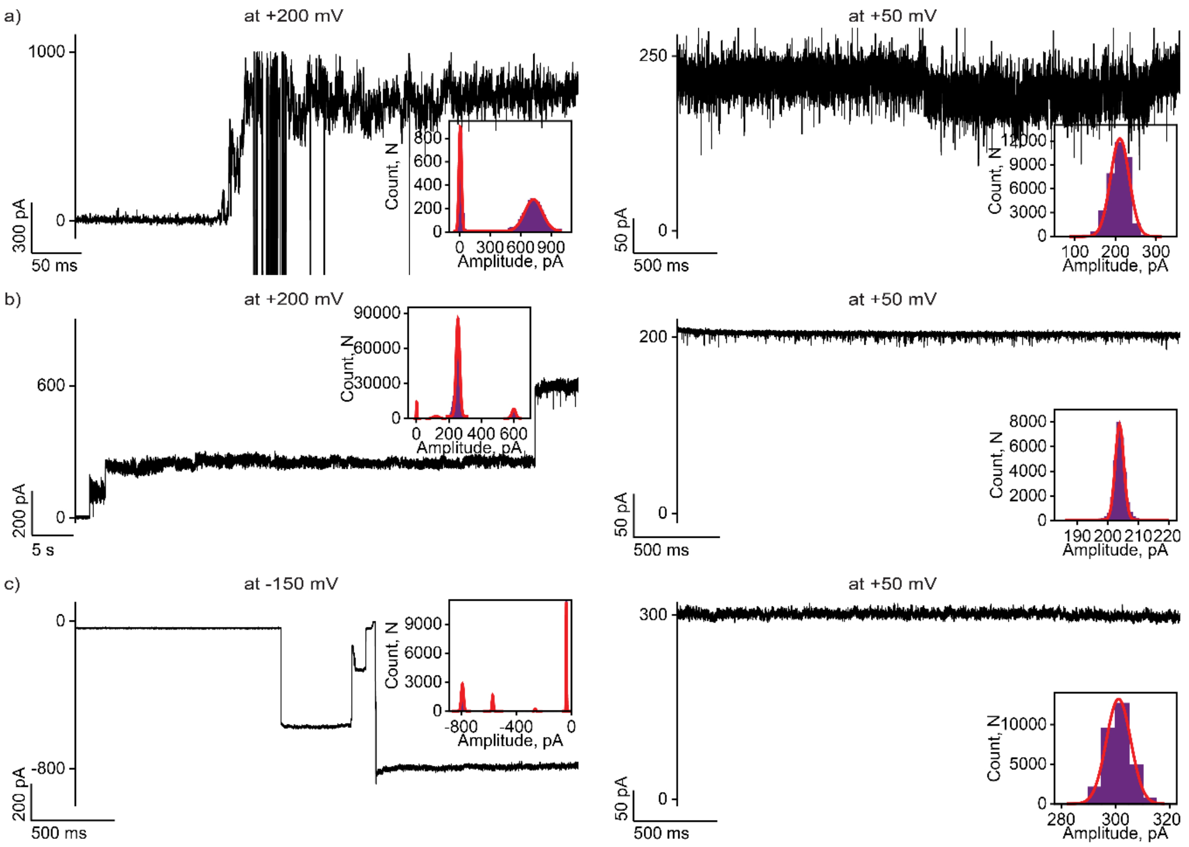


**Figure S2: Real-time insertions and large pore activity of pPorU pores in planar lipid bilayers**

**a)** Electrical recording of single pPorU channel insertion into DPhPC lipid bilayer at +200 mV and the same channel at +50 mV. **b)** Electrical recording of single pPorU channel insertion into DPhPC lipid bilayer at +200 mV and the same channel at +50 mV. **c)** Electrical recording of single pPorU channel insertion into DPhPC lipid bilayer at -150 mV and the same channel at +50 mV. Inset shows the corresponding all-points amplitude histogram. The current signals were filtered at 2 kHz and sampled at 10 kHz. Electrolyte: 1 M KCl, 10 mM HEPES, pH 7.4.

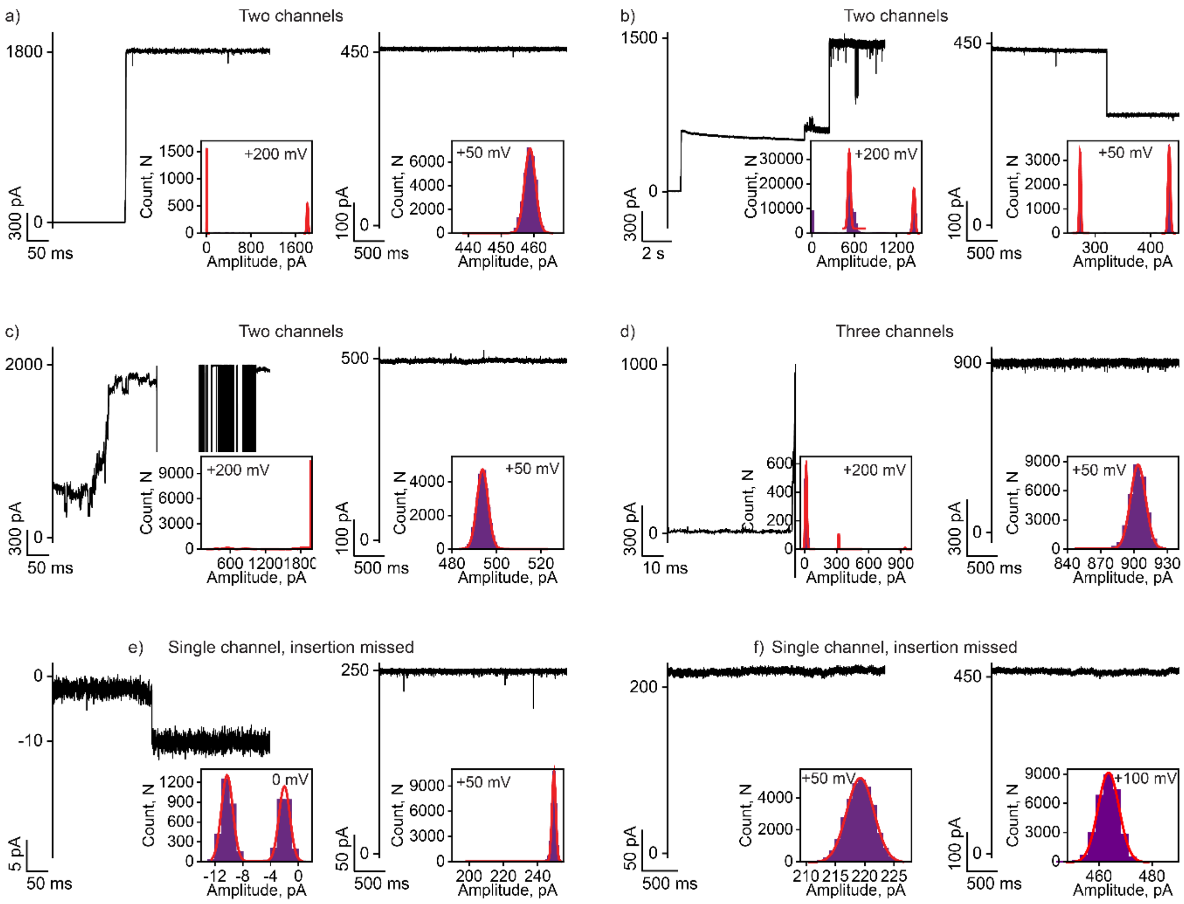


**Figure S3: Large stable pPorU pores in planar lipid bilayers at different conditions**

a), b), c) Electrical recording of two pPorU channels inserting into DPhPC lipid bilayer at +200 mV and the same channels at +50 mV. d) Electrical recording of three pPorU channels inserting into DPhPC lipid bilayer at +200 mV and the same channels at +50 mV. e) Electrical recording of single pPorU channel inserting into DPhPC lipid bilayer at +0 mV and the same channel at +50 mV. f) Electrical recording of single pPorU channel in the DPhPC lipid bilayer at +50 mV and +100 mV. The insertion of this channel was missed. Inset shows the corresponding all-points amplitude histogram. The current signals were filtered at 2 kHz and sampled at 10 kHz. Electrolyte: 1 M KCl, 10 mM HEPES, pH 7.4.

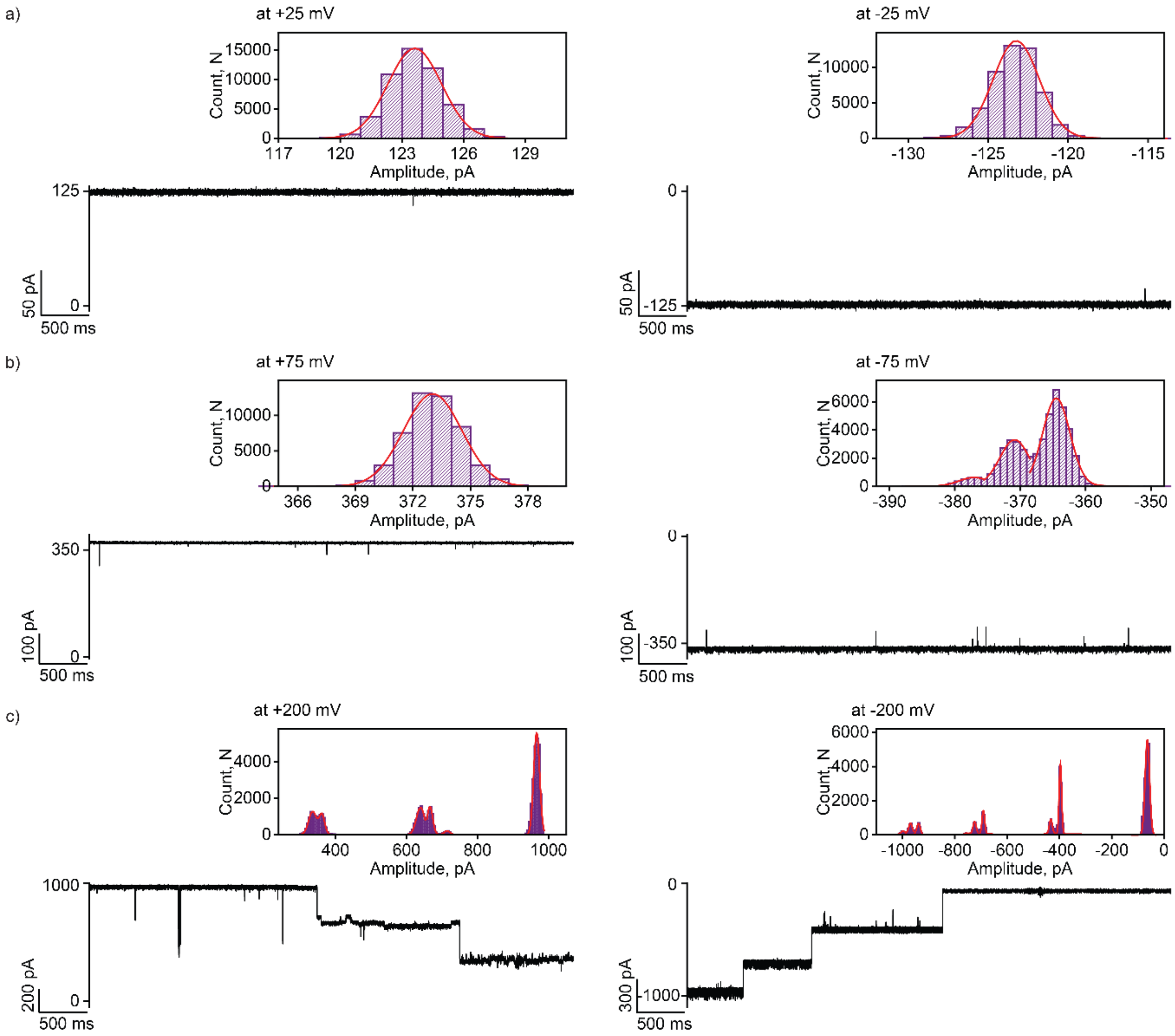


**Figure S4: Gating of large pPorU pores at different voltages**

a) Electrical recording of pPorU channel in DPhPC lipid bilayer at +25 mV and -25 mV. b) Electrical recording of pPorU channel in DPhPC lipid bilayer at +75 mV and -75 mV. c) Electrical recording of pPorU channel in DPhPC lipid bilayer at +200 mV and -200 mV. Inset shows the corresponding all-points amplitude histogram. The current signals were filtered at 2 kHz and sampled at 10 kHz. Electrolyte: 1 M KCl, 10 mM HEPES, pH 7.4.

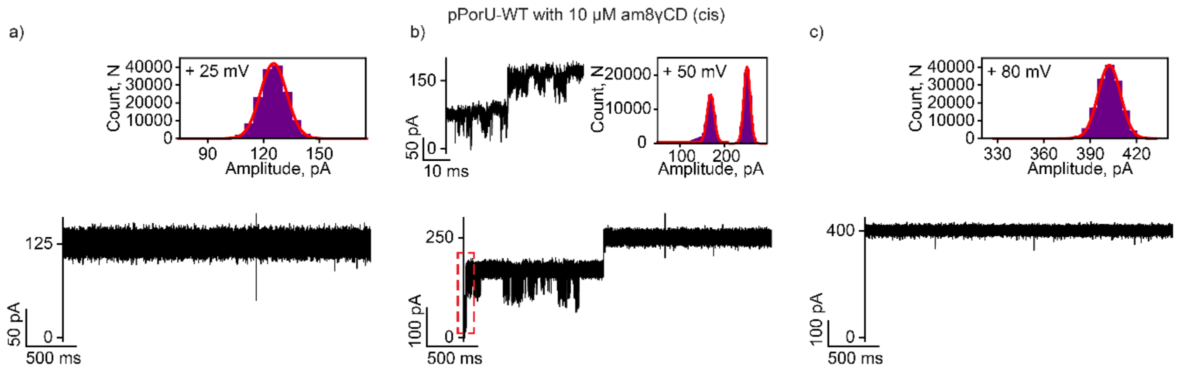


**Figure S5: Interaction of cationic Gamma CD with large pPorU pores**

Electrical recordings showing the interaction of cationic gamma-cyclodextrin (am$_8$γCD) with single pPorU pore (10 µM, cis) at a) +25 mV b) +50 mV and c) +80 mV against electrophoretic mobility with clear release events. Inset shows the corresponding all-points amplitude histogram and ion current recordings at an expanded time scale. The current signals were filtered at 10 kHz and sampled at 50 kHz. Electrolyte: 1 M KCl, 10 mM HEPES, pH 7.4.

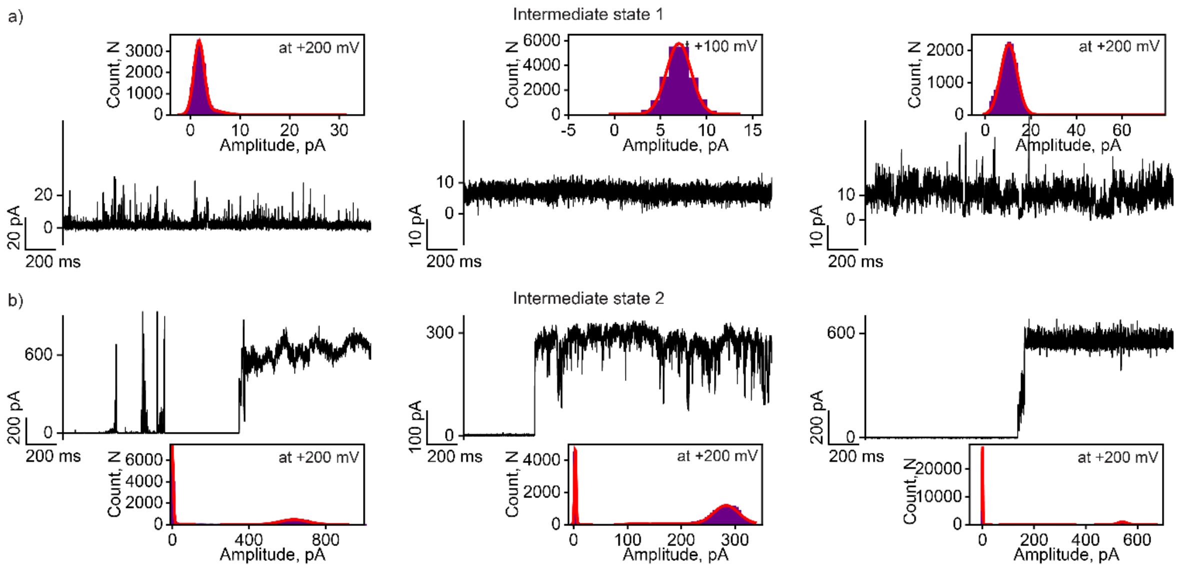


**Figure S6: Intermediate states of pPorU pores formed at lower voltages**

a) Electrical recording of intermediate conductance states at +100 mV and +200 mV producing low current. b) Electrical recording of different intermediate states at +200 mV showing higher current. Inset shows the corresponding all-points amplitude histogram. The current signals were filtered at 2 kHz and sampled at 10 kHz. Electrolyte: 1 M KCl, 10 mM HEPES, pH 7.4.

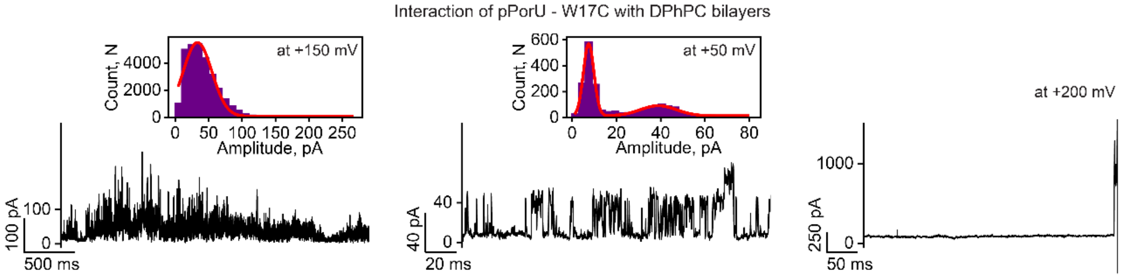


**Figure S7: Electrical recordings of pPorU-W17C peptides in planar lipid bilayers**

Electrical recording of the interaction of pPorU-W17C peptides with DPhPC lipid bilayer showing fluctuating interaction at +150 mV. Electrical recording showing the interaction of the pPorU-W17C peptides with DPhPC lipid bilayer showing two states at +50 mV. Electrical recording of the interaction of pPorU-W17C peptides with the DPhPC lipid bilayer showing the disruption of the bilayer at +200 mV. Inset shows the corresponding all-points amplitude histogram. The current signals were filtered at 2 kHz and sampled at 10 kHz. Electrolyte: 1 M KCl, 10 mM HEPES, pH 7.4.

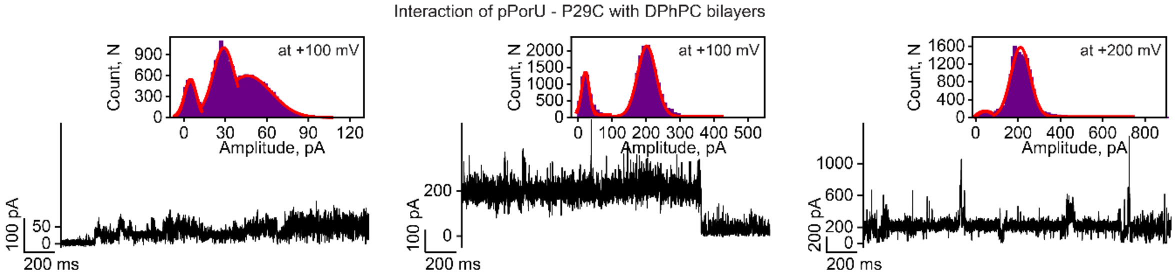


**Figure S8: Electrical recordings of pPorU-P29C peptides in planar lipid bilayers**

Electrical recordings of pPorU-P29C channel activity in DPhPC lipid bilayer showing conductance similar to intermediate steps at +100mV and +200 mV. Inset shows the corresponding all-points amplitude histogram. The current signals were filtered at 2 kHz and sampled at 10 kHz. Electrolyte: 1 M KCl, 10 mM HEPES, pH 7.4.

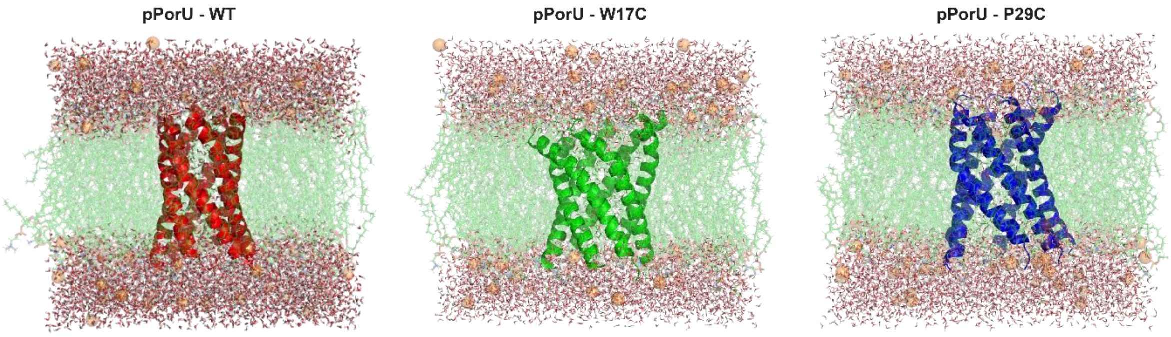


**Figure S9: MD simulations of pPorU-WT, pPorU-W17C and pPorU-P29C**

Simulated hexapeptide pore structures for the native pPorU structure (red), as well as the pPorU-W17C (green) and pPorU-P29C (blue) mutants.

Using the on-line Charmm GUI, each resulting pore was embedded into a bilayer composed of POPC, POPE and POPG lipids in a 3:1:1 ratio, with cytoplasmic and extracellular buffers (extending 15.0 Å above and below the bilayer) composed of water, 67 K+ ions and 21 Cl- ions. Extensive restrained constant pressure structural refinements were carried out in NAMD, representing the structures via the Charmm 3.6 force field. Given a thermostatic temperature of 303.15K, an initial 10,000 step relaxation was performed using quadratic positional restraints on all protein backbone atoms, via a restraint scaling factor of 10.0. A second 12,500 step restraint relaxation was then conducted on the resulting system, using a scaling factor of 5.0. Four further relaxations were then conducted (of respective lengths 12,500, 25,000, 25,000, and 25,000 steps) with restraint scaling factors diminishing according to the progression 2.5, 1.0, 0.5 and 0.1). A final production simulation was then run for 4.0 nS, at constant pressure, with no positional restraints

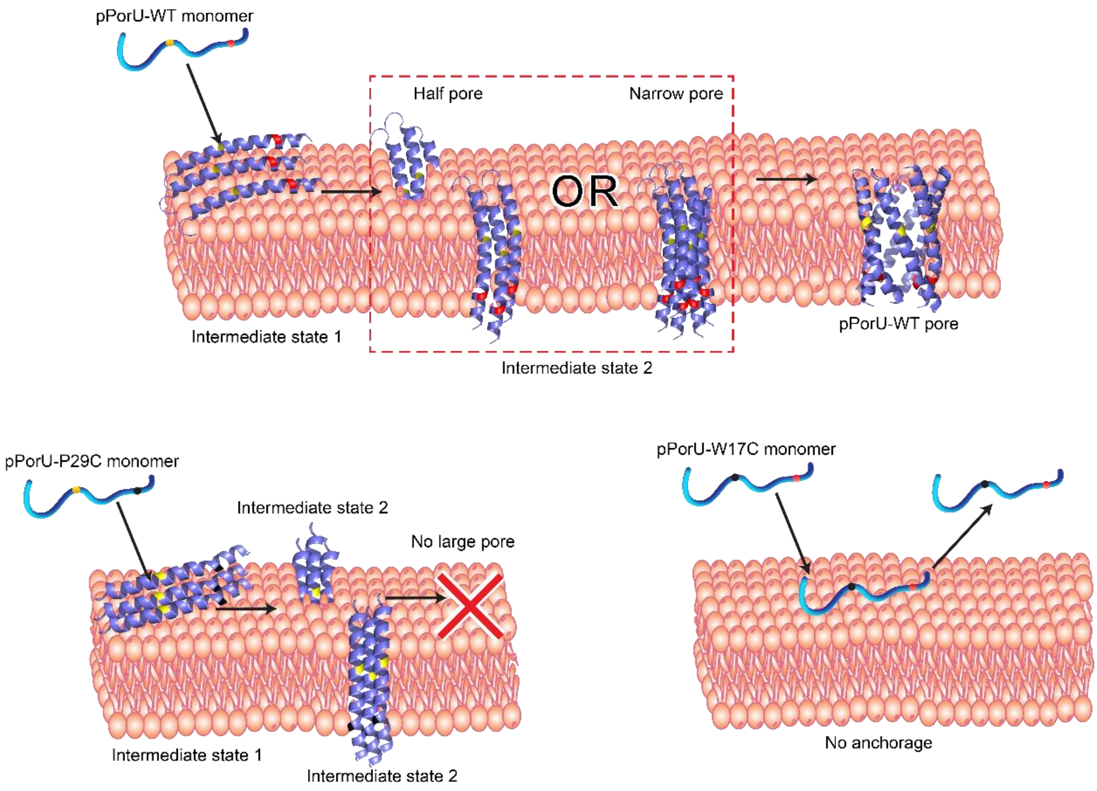


**Figure S10: Model showing membrane insertion and pore formation of pPorU peptides**

Proposed model showing the different steps in the assembly of the large pore, including intermediate step 1, intermediate step 2 and pore formation. The model also shows the impact of mutation of critical residues on the assembly pathway.
